# Testing differential effects of periodicity and predictability in auditory rhythmic cueing of concurrent speech

**DOI:** 10.64898/2026.03.11.711109

**Authors:** Jessica MacLean, Mengyuan Zhou, Gavin M. Bidelman

## Abstract

Entrainment and predictive coding aid speech perception in both quiet and noisy environments. Isochronous, periodic auditory rhythmic cues facilitate entrainment and temporal expectations which can benefit encoding and perception of target speech. However, most studies using isochronous cues confound periodicity with predictability. To this end, we characterized how systematic changes in the acoustic dimensions of stimulus rate, target phase, periodicity, and predictably of an entraining sound precursor impact the subsequent identification of concurrent speech targets. Target concurrent vowel pairs were preceded by rhythmic woodblock cues which were either periodic-predictable (PP, isochronous rhythm), aperiodic-predictable (AP, accelerating rhythm), or aperiodic-unpredictable (AU, random rhythm). The number of pulses per rhythm was roved to further manipulate predictability. Stimuli also varied in presentation rate (2.5, 4.5, 6.5 Hz) and target speech phase (in-phase, 0°; out-of-phase, 90°, 180°) relative to the preceding entraining rhythm. We also measured participants’ musical pulse continuation and standardized speech-in-noise perception abilities. We did not observe any effects of stimulus rhythm, rate, or target phase on target speech identification accuracy. However, reaction times were slowest at the nominal speech rate (4.5 Hz) and were most disrupted by out-of-phase presentations following the PP rhythm. Double-vowel task performance was associated with stronger musical pulse continuation abilities, but not speech-in-noise perception. Our results support the notion that entraining rhythmic cues rely on top-down processing but are relatively muted when stimulus predictability is unknown. Additionally, we find that individual differences in musical pulse perception may underlie the benefits of rhythmic cueing on subsequent speech perception.

## 1 Introduction

Temporal context is highly important to speech perception. Listeners use natural speech rhythms to facilitate comprehension in various ways, including chunking speech into meaningful units, predicting subsequent temporal patterns, and separating target speech from background noise (Ding et al., 2014; McAuley et al., 2020; Smith et al., 2024). This ability is likely supported by neural entrainment, or the yoking of ongoing low-frequency neural oscillations to periodicities in external stimuli such as speech (Ding and Simon, 2014). Neural entrainment to speech underlies successful comprehension across listening environments (Peelle and Davis, 2012; Riecke et al., 2018; Etard and Reichenbach, 2019). As such, an emerging body of literature has investigated the beneficial effects of preceding rhythmic cues on neural entrainment to, and later perception of, speech material.

“Forward entrainment” refers to the phenomenon that when neural oscillations couple with an external stimulus, the oscillations continue for a brief window (up to a few cycles) after the stimulus stops (Lakatos et al., 2019; Saberi and Hickok, 2022; 2023). This effect enhances perceptual sensitivity in tone detection and pitch discrimination in noise (Jones et al., 2002; Hickok et al., 2015; Farahbod et al., 2020; Solli et al., 2024) likely through oscillatory coupling and attentional direction to salient points in time (i.e., Dynamic Attending Theory; Jones and Boltz, 1989; Large and Jones, 1999). Similarly, rhythmic cues have been shown to benefit speech processing in quiet and noise (Falk et al., 2017; te Rietmolen et al., 2025), but there have been few systematic investigations of how rhythmic cue parameters influence speech perception.

Most studies utilize a metronome-like cue that is perfectly isochronous (Jones et al., 2002; te Rietmolen et al., 2025). This potentially confounds cue periodicity and predictability in enhancing speech perception. In attempts to address this question, Solli et al. (2024) presented periodic, predictable (PP, metronome-like), aperiodic, yet predictable (AP, increase in speed at a steady inter-stimulus interval ratio), and aperiodic, unpredictable (AU, random) cues prior to tone probes in a pitch discrimination in noise task. While both PP and AP cues enhanced perception of the target tone, only PP cues facilitated neural entrainment. While this study demonstrated that periodicity provides additional benefits over predictability to perception, the rhythmic stimuli always contained the same number of entraining pulses. Unfortunately, this stimulus design does not allow one to tease apart the relative benefits of periodicity vs. predictability on task facilitation; targets were always, to some degree, highly predictable following a fixed number of events.

Neural entrainment to speech seems to be enhanced at certain rates. In particular, the neural tracking of speech is enhanced at 4-5 Hz (He et al., 2023; 2024; Momtaz and Bidelman, 2024), the purported “ideal” syllable rate observed across many of the world’s languages (Assaneo and Poeppel, 2018; Assaneo et al., 2019). Some rhythmic cueing studies have shown improvements to speech-in-noise comprehension when cued at both the syllable rate and slower, “foot” rates (te Rietmolen et al., 2025). Thus, rhythms may facilitate speech processing differently depending on their speed.

Speech itself is also quasi-periodic, meaning that neural entrainment to speech could be more robust to slight differences in target timing. Indeed, Solli et al. (2024) found that PP cues elicited phase-specific benefits to target pitch perception when the target tone was presented in the expected phase (0°) relative to the cue interstimulus interval (ISI), consistent with predictions of Dynamic Attending Theory (Jones and Boltz, 1989; Large and Jones, 1999). However, for AP cues, this phase-specificity was not observed. Thus, while alteration of target speech phase might enhance or disrupt rhythmic cue benefits to later auditory perception, such effects seem heavily tied to the predictability of the upcoming stimulus.

Here, we extend findings of Solli et al. (2024) by presenting similarly constructed (PP, AP, and AU) cues prior to *speech* targets to test how preceding rhythm periodicity and predictability impact later speech perception. We selected double-vowel (DV) mixtures (Assmann and Summerfield, 1989; 1990) as the speech target as these stimuli have been used widely in auditory perception studies to investigate influences of short- and long-term auditory experiences on the rapid perception of concurrent speech without observance of ceiling effects (Alain et al., 2007; Yellamsetty and Bidelman, 2018; 2019; MacLean et al., 2024; MacLean et al., 2025). To parametrically test speed effects, we varied the rate of rhythms between 2.5, 4.5, and 6.5 Hz. To test the effects of cued-target phase, we presented DV targets in phase (0°) or out of phase (-90° early, 180° late) relative to preceding rhythms. Critically, we also roved the number of pulses in the preceding rhythmic cue to prevent listeners from merely anticipating the speech signal after a fixed number of events. This has not been well controlled in prior studies (e.g., Hickok et al., 2015; Solli et al., 2024). Collectively, our design attempted to disentangle the effects of stimulus periodicity (bottom-up neural entrainment) and predictability (top-down predictive coding) on target speech identification.

## 2 Materials and Methods

### 2.1 Participants

Twenty-three young adults (ages 18-35 years; mean ± SD: 22.04 ± 3.30, 16 female) participated in this study. All participants had bilateral normal hearing thresholds (<25 dB HL) at octave frequencies between 250 and 8000 Hz, were fluent in American English, and reported no history of psychiatric or neurologic disorders. Participants had varied amounts of music training (range: 0-23 years; mean ± *SD*: 7.83 ± 7.06 years). Handedness was assessed through the Edinburgh Handedness Inventory (range: -15 to 100%; mean ± SD: 64.79 ± 32.98%) (Oldfield, 1971). All study procedures, including written informed consent, were performed in accordance with a protocol approved by the Indiana University Institutional Review Board (#23256).

### 2.2 Stimuli and task

#### 2.2.1 Task overview

On each trial, target speech was preceded by one of three rhythmic cues [periodic-predictable (PP), aperiodic-predictable (AP), aperiodic-unpredictable (AU)] presented at a rate of 2.5, 4.5, or 6.5 Hz (described below; **Figure 1A**). Targets could also occur in- or out-of-phase with respect to the preceding rhythm. Trials were blocked by rate; rhythmic cues and phase were ordered randomly within rate. There were 24 trials per condition resulting in a total of 648 trials in the task (=3 rates x 3 phases x 3 rhythms x 24 repetitions). We used a Latin square to randomize block order across participants. On each trial, participants identified the targets (double vowels) via keyboard press. Accuracy (percent correct identification of both vowels) and reaction time (RT) were measured. Auditory stimuli were presented binaurally at 79 dB SPL through ER-2 insert earphones (Etymotic Research, Elk Grove, IL) via a MATLAB-controlled TDT RZ6 interface (Tucker-Davis Technologies, Alachua, FL).

**Figure 1.**
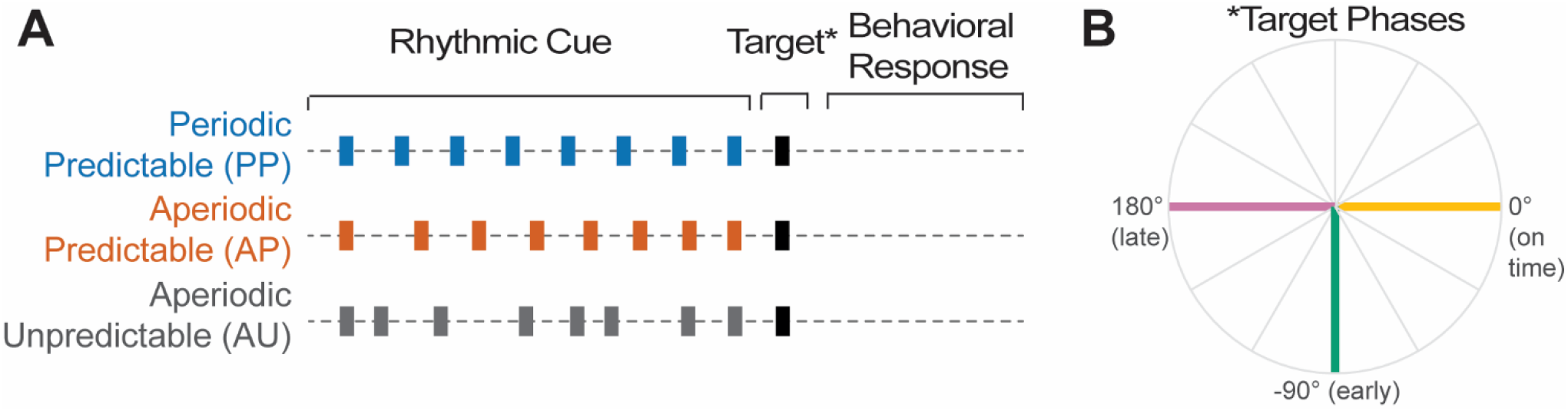
Stimulus design to assess rhythmic cueing benefits on speech perception. **(A)** Target double vowel pairs (*) were preceded by one of three rhythmic cue conditions: Periodic Predictable—PP (isochronous rhythm), Aperiodic Predictable—AP (accelerating rhythm), and Aperiodic Unpredictable—AU (random rhythm). Rhythms were presented at one of three nominal rates (2.5, 4.5, 6.5 Hz). Each rhythm consisted of a train of woodblock sounds. **(B)** Targets (i.e., vowel pairs) also occurred in- or out-of-phase (lead, lag) relative to the preceding rhythm.

#### 2.2.2 Speech targets

Target stimuli consisted of double vowel pairs (MacLean et al., 2024; MacLean et al., 2025). Each pair consisted of two, steady-state vowels (/a/, /e/, and /i/) presented simultaneously in three unique combinations (i.e., /a/ + /e/; /e/ + /i/; /a/ + /i/). Vowels were never paired with themselves. Individual vowel sounds were created with a Klatt-like synthesizer (Klatt, 1980) via MATLAB v2021 (The MathWorks, Inc., Natick, MA). Each was 100 ms in duration with 10-ms cos^2^ onset/offset ramping to prevent spectral splatter. Vowel pairs contained one vowel with a fundamental frequency (F0) of 150 Hz and another of 190 Hz (4 semitones). F0 and the first two formant frequencies (*F1*_a,e,i_ = 787, 583, 300 Hz; *F2* _a,e,i_ = 1307, 1753, 2805 Hz) remained constant for the entire stimulus duration. Prior to the main task, we required all participants to identify single vowels with 100% accuracy to ensure that task performance would measure concurrent speech identification rather than isolated sound labeling. The actual experimental task required participants to identify *both* vowels in a pair for a correct response.

#### 2.2.3 Rhythmic trains

Each rhythm was designed to manipulate the periodicity and/or predictability of the preceding rhythmic cue on subsequent target speech detection. Rhythms were constructed from a train of 100 ms woodblock cues (sampled from Flat music notation software, Tutteo, Inc., Claymont, DE). Rhythms were presented at three different nominal rates (2.5, 4.5, 6.5 Hz; separate blocks) to assess the effects of stimulus speed on inducing entrainment. Each rhythmic condition contained 7-9 woodblock sounds, roved across trials to prevent participants from anticipating the impending double-vowel target. For a given rate, there were three types of rhythmic cues preceding the target (**Fig. 1A**), modeled after previous studies designed to distinguish effects of stimulus periodicity and predictability on auditory perception (Solli et al., 2024). Participants were instructed to use preceding auditory rhythms to help them predict subsequent vowel pairs.

#### 2.2.4 Periodicity/predictability manipulation

For a particular rate, periodic-predictable (PP) rhythms consisted of a train of steady inter-stimulus intervals (ISIs) at the same period (i.e., isochronous metronome) based on the stimulus rate of that block. Aperiodic-predictable (AP) rhythms started at 1.5x slower than the nominal rate with pulse ISI accelerating (on a log scale) to reach the nominal rate by the end of the train. AP stimuli removed stimulus periodicity but retained predictability (Solli et al., 2024). Aperiodic-unpredictable (AU) rhythms were pseudo-randomly jittered versions of the PP cues with the ISI jittered (0.5-1.5x) around the nominal rate. AU stimuli lacked both periodicity and predictability and were thus expected to be uninformative for target perception. Regardless of condition, the time between the entire rhythmic train and double vowel target was always identical (i.e., the nominal ISI for a given rate) to allow for phase manipulation of the target.

#### 2.2.5 Phase manipulation

In addition to rate and rhythmic periodicity/predictability, we varied the relative phase of targets with respect to the preceding rhythm to further examine the putative effects of entrainment on speech perception. Target vowel pairs were presented in three phases relative to the final pulse in the preceding rhythmic train (**Figure 1B**): in phase (0°), out of phase early (-90° lead), or out of phase late (180° lag) relative to the preceding ISI.

### 2.3 Speech-in-noise perception

In addition to improving detection of auditory targets, entrainment has also been linked with improved SIN perception (Riecke et al., 2018; Vanthornhout et al., 2018; Etard and Reichenbach, 2019). To assess relationships between our entertainment and SIN performance, we administered two lists from the Quick Speech-in-Noise (QuickSIN) test (Killion et al., 2004). Participants were asked to repeat low-context target sentences presented in increasing levels of four-talker babble. The outcome of QuickSIN is a clinically-normed measure of “SNR loss,” reflecting the dB SNR for 50% keyboard recall performance (0-3 dB SNR loss is considered normal for this task).

### 2.4 Musical beat perception

The Beat-Drop Alignment Test (BDAT; Cinelyte et al., 2022) was used to measure participants’ musical pulse continuation abilities and provide an external measure of entrainment abilities. In the BDAT, each trial uses naturalistic musical stimuli to establish a beat, which then “drops out” (i.e., stops) and is followed by a single probe tone. Participants indicate whether the probe tone falls “on” or “off” the beat relative to the musical rhythm established just prior. For “off” condition stimuli, probe tones are displaced by 7 equal step sizes between 15% and 45% of the beat period. Displacements occur before and after “on” beats. The test is adaptive and results in a z-score indicating participants’ perceptual pulse continuation threshold for entraining to the beat (higher scores indicate better performance). Critically, no concurrent cues are presented during judgments of the probe. Thus, to perform well on the BDAT, listeners must form a mental representation of the rhythmic beat structure and later compare this internal representation to the single probe event.

### 2.5 Statistical analysis

We utilized generalized linear mixed-models in R (version 4.2.2, lme4 package) to analyze the dependent variables. All models utilized fixed predictors of rhythm condition (3 levels: PP, AP, AU), rate (3 levels: 2.5, 4.5, 6.5 Hz), and phase (3 levels: 0, -90, 180°), their full two- and three-way interactions, and scaled BDAT scores as a continuous covariate. Models included both random intercepts for subject and random slopes for rate as allowed by model convergence (Barr et al., 2013).

Trial level accuracy data were coded binarily (correct= 1, incorrect= 0) and fit to a logistic model with glmer using a binomial link function [i.e., logit(*corr) ∼phase*rate*cond + BDAT + (1 + rate*|*sub)*]. Wald chi-square tests (type III) were used to evaluate significance for glmer models.

For RTs, only correct trials were analyzed. RTs were also adjusted to account for the ISI by removing a duration of 1/rate from each RT value and were then log-transformed to normalize the data. We used lmer models to analyze the continuous RT data [i.e., *RT ∼ phase*rate*cond + BDAT + (1 + rate*|*sub)*].

Pairwise comparisons were adjusted using Tukey corrections. Effect sizes are reported as odds ratio (OR) or partial eta squared 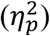 for logit and Gaussian models, respectively. Degrees of freedom were calculated using Satterthwaite’s method. We performed partial correlation analyses (ppcor package) between double vowel identification accuracy and RT (averaged across conditions), external measures (QuickSIN, BDAT), and demographics (e.g., music training) to assess bivariate relationships within our dataset while controlling for influences from other variables.

## 3 Results

### 3.1 Speech identification: Accuracy (%)

Trial-level accuracy was not affected by target phase (*X*^2^(2) = 2.26, *p* = 0.32), rate (*X*^2^(2) = 0.46, *p* = 0.79), rhythm condition (*X*^2^(2) = 0.86, *p* = 0.65), nor their interactions (all *p* > 0.30; **Figure 2A**). However, we found that BDAT scores positively predicted task accuracy (*X*^2^(1) = 5.72, *p* = 0.017, OR = 1.84); better musical pulse continuation scores predicted greater double-vowel identification after rhythmic cueing (**Figure 2B**).

**Figure 2.**
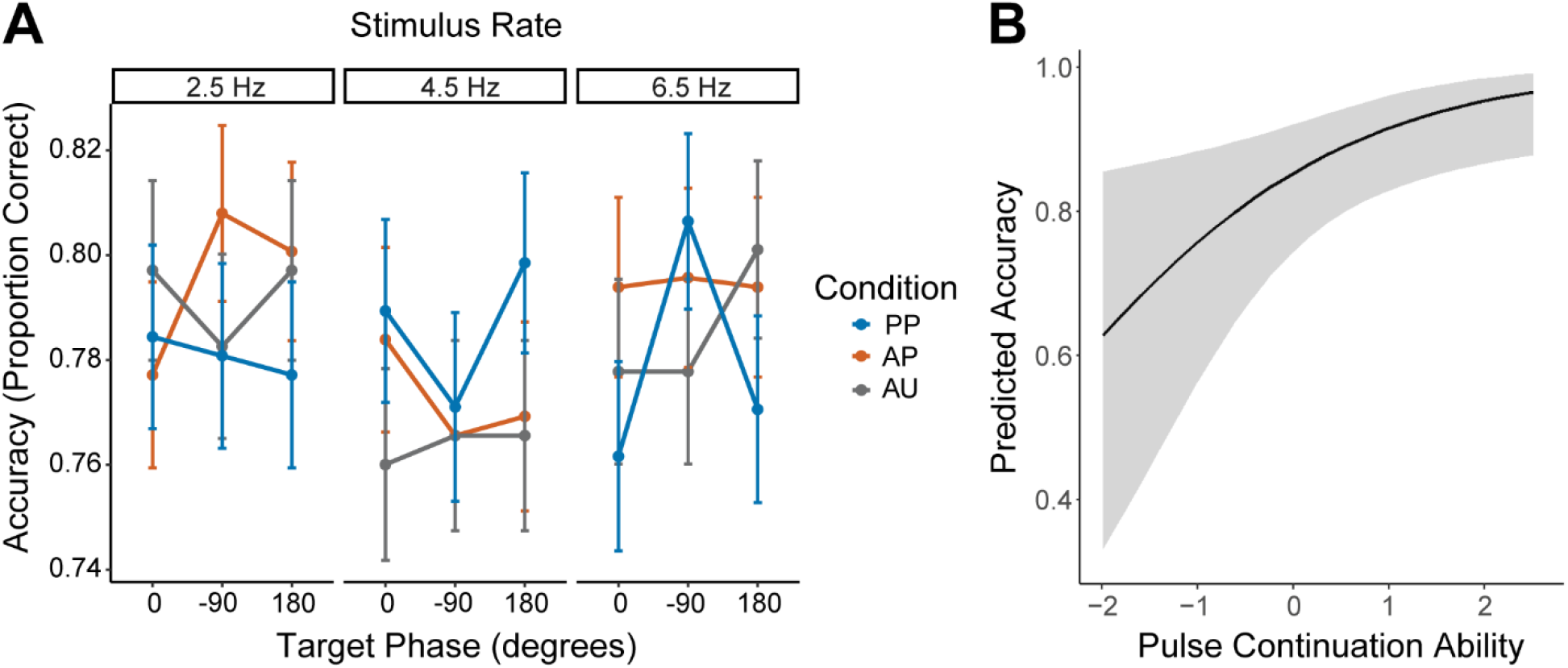
Speech identification accuracy across experimental manipulations and relation to musical pulse continuation scores. **(A)** Accuracy in the primary task was invariant to experimental manipulations of rate, periodicity, predictability, and phase. **(B)** Higher BDAT scores predicted overall more accurate double-vowel speech perception in the rhythmic cueing task. Line = logistic regression model fit. Shading: 95% CI. Error bars: + 1 S.E.M. PP, periodic-predictable; AP, aperiodic-predictable, AU, aperiodic-unpredictable.

### 3.2 Speech identification: Reaction times

RTs for speech identification are shown in **Figure 3**. RTs were modulated by a target phase x rhythm type interaction (*F*(11587) = 2.786, *p* = 0.0250) due to longer RTs for the -90° relative to the 180° (*p* = 0.0031) and 0° phase (*p* = 0.0667) target positions, particularly in the PP rhythm condition (**Fig. 3A**). RTs were also modulated by the rate of the preceding rhythm (*F*(21.5) = 6.99, *p* = 0.0045), largely due to longer RTs for the 4.5 Hz relative to the 2.5 Hz stimulus rate (*p* = 0.0006; **Fig. 3B**). No other main effects or interactions were observed (all *p* > 0.1).

**Figure 3.**
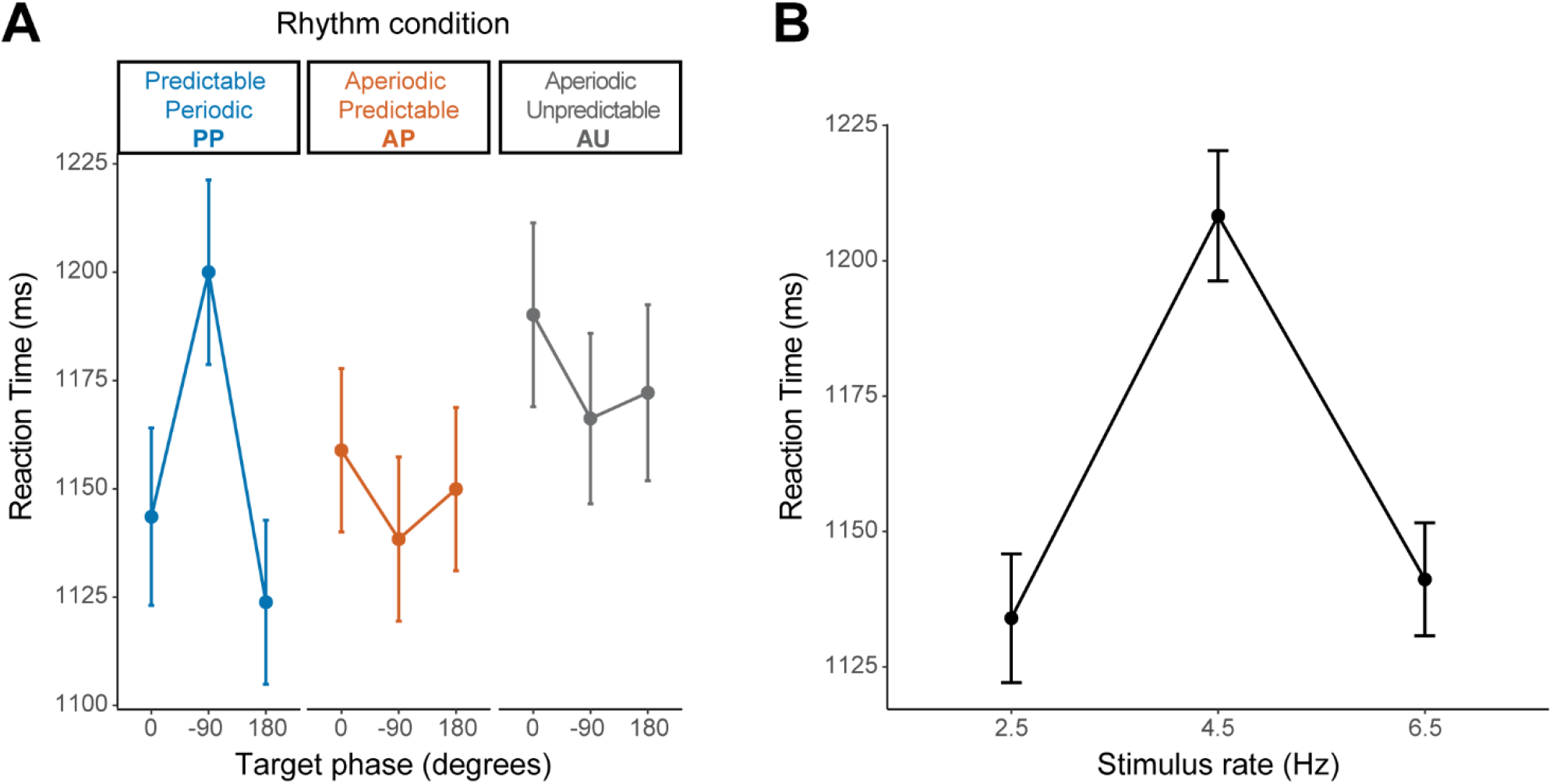
Reaction times for speech identification following entraining rhythms. **(A)** RTs speed for identifying double-vowels when positioned at different phases following (a)periodic/periodic and (un)predictable rhythms. There was an interaction between phase and rhythm with slower RTs for the -900 phase targets. **(B)** Main effect of rate on speech identification speed. Slower RTs were observed for the 4.5 Hz rate. Error bars: + 1 S.E.M.

### 3.3 Correlations between entrainment measures, SIN, and music demographics

We performed partial correlations between speech perception accuracy (collapsed across task conditions), SIN performance (QuickSIN), musical pulse continuation ability (BDAT), and years of music training (**Figure 4**). We observed a significant relationship between years of music training and BDAT scores (**Fig. 4A**; *R* = 0.533, *p* = 0.016), replicating prior findings (Cinelyte et al., 2022). QuickSIN SNR loss scores did not correlate with task accuracy (**Fig. 4B;** *R* = -0.18, *p* = 0.448), RT (**Fig. 4C;** *R* = 0.252, *p* = 0.284), or BDAT scores (**Fig. 4D**; *R* = 0.256, *p* = 0.275). All other relationships in the partial correlation matrix were nonsignificant (data not shown; all *p*s > 0.05).

**Figure 4.**
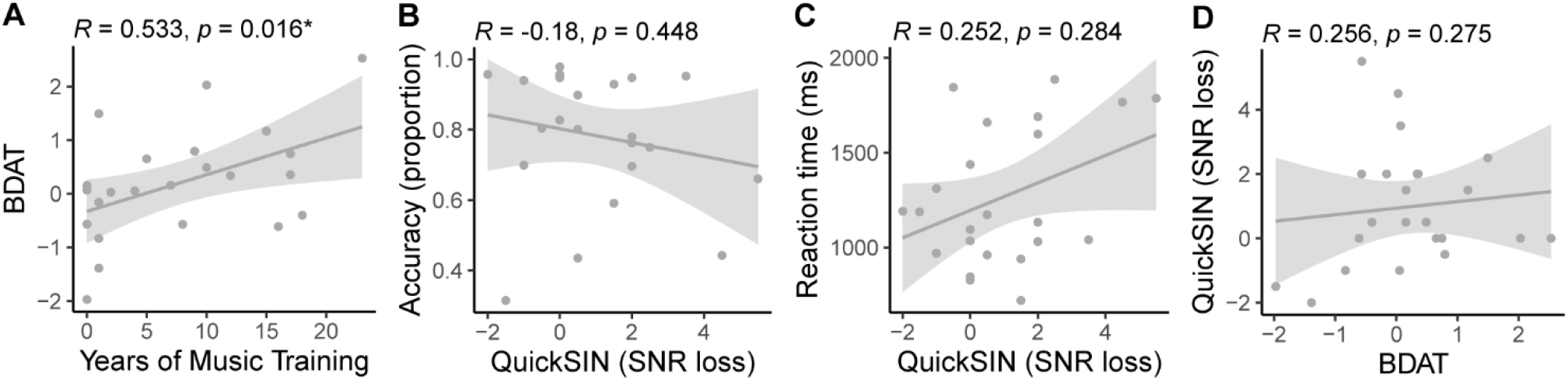
Correlations between task accuracy and external measures. Statistics show the partial correlations between each bivariate pair adjusted for the other variables. **(A)** Years of music training correlated with BDAT scores. **(B-C)** QuickSIN scores did not correlate with task accuracy, RTs, nor BDAT scores. Shading: 95% CI, *p < 0.05.

## 4 Discussion

Through manipulation of rate, periodicity, and target phase in a rhythmically-cued concurrent vowel paradigm, we aimed to assess the effects of preceding musical rhythms (inducing perceptual entrainment) on subsequent speech perception. We found that: a) concurrent vowel identification speed (but not accuracy) was impacted by preceding music rhythms; b) external measures of musical pulse continuation abilities positively correlated with task accuracy and related to listeners’ amount of music training; and c) cued speech perception did not relate to external measures of speech-in-noise (SIN) perception.

### 4.1 No impact of rhythmic entrainment on speech perception accuracy

We found that concurrent vowel identification *accuracy* was not influenced by rate, rhythm (a)periodicity/predictability, or target phase manipulations. Though prior rhythmic cues have been shown to influence later auditory detection and speech perception (Falk et al., 2017; Berthault et al., 2024; te Rietmolen et al., 2025), this effect is not universally observed (Bauer et al., 2015; Lin et al., 2022; Román-Caballero et al., 2024). Possible reasons that our stimulus manipulations did not elicit perceptual accuracy benefits include a) roving of the rhythmic cue, b) possible lack of attention to rhythmic cues, and c) lack of titrated stimulus difficulty.

#### 4.1.1 Roving of rhythmic cue

Though we had anticipated changes in speech identification scores, the lack of accuracy effects here provides novel information about the limits of forward entrainment. Most studies have used a fixed number of entraining sounds when exploring entrainment effects (Jones et al., 2002; Hickok et al., 2015; Farahbod et al., 2020). Here, we roved the entraining rhythm, so that the number of sounds varied from trial to trial. This retained cue periodicities but prevented participants from counting down the number of sounds in the less predictable stimulus condition (AU). As a result, our stimuli isolated the effects of pure periodicity/predictability, rather than building anticipatory patterns which participants could learn across the experiment. Our results suggest that roving mutes the benefits of rhythmic cue entrainment previously seen using similar cues (Bauer et al., 2015; Farahbod et al., 2020; Solli et al., 2024). Because roving likely disrupts cue predictability, previous benefits seen from rhythmic cues may rely more heavily on top-down anticipatory effects on perception rather than bottom-up entrainment, per se.

Alternatively, altering the number of entraining sounds may disrupt nesting of the cued faster rate (here, 4.5 Hz) with slower rates in the oscillatory hierarchy. Other studies have shown perceptual cueing benefits for coupled delta-beta and delta-theta rhythms (Chang et al., 2019; Kubetschek and Kayser, 2021), which would be disrupted through roving the number of pulses in rhythm sequences. At the very least, our data argue that roving is a crucial addition to future forward entrainment experiments to disentangle periodicity/predictability from trivial anticipation effects that could equally underly the auditory perceptual facilitation from rhythmic cueing.

#### 4.1.2 Attention to rhythmic cue

The persistence of neural entrainment following a rhythmic stimulus is fleeting, lasting for a few cycles at maximum (L’Hermite and Zoefel, 2023; Saberi and Hickok, 2023). Importantly, this effect may only be observed under active attention to the entraining stimulus (Lakatos et al., 2013; Farahbod et al., 2020; Bouwer, 2022). Without strong attentional capture to the preceding rhythms, participants may not experience attention-related enhancements in neural entrainment to the rhythms, tempering benefits to concurrent vowel perception. Because we roved our stimuli, it is possible that participants partially ignored the woodblock cues, which would decrease the cognitive dynamic attending benefits of a rhythmic cue (Jones and Boltz, 1989; Large and Jones, 1999). Still, we find the trivial account that participants simply “tuned out” the rhythm cue unlikely, since RT decision speeds did vary with the preceding rhythmic rate. This confirms that rhythmic priming affected subsequent speech perception, albeit in a weak manner.

#### 4.1.3 Titrated performance

Contrary to our hypothesis, the additional predictability and periodicity in the rhythm cue afforded by the AP and PP conditions, respectively, did not benefit concurrent vowel identification. This contrasts with prior work utilizing similar rhythmic cues for pitch identification (Solli et al., 2024). However, Solli et al. presented target tones in noise and measured individualized SNR thresholds to equate performance across listeners at 75% accuracy. In contrast, we used double vowel mixtures presented in quiet and did not titrate performance. However, we note that listeners in our task achieved a roughly similar level of performance (75-80%). The speech-on-speech nature of our double-vowel stimuli aside, studies suggest that entraining rhythm cues may be most helpful in difficult or noisy listening environments (McAuley et al., 2020; Pearson et al., 2023) or when there is level-uncertainty in near-threshold detection (Farahbod et al., 2020). Thus, the lack of noise or tailored difficulty in our task may also explain the lack of strong rhythmicity effect on speech perception accuracy.

### 4.2 Rhythmic entrainment impacts the speed of speech identification decisions

Unlike accuracy, *decision speeds* for target speech identification were modulated by the rate of priming rhythms. We had expected faster RTs at 4.5 Hz due to selective enhancement of neural entrainment at this rate (Assaneo and Poeppel, 2018; He et al., 2023; Momtaz and Bidelman, 2024). However, our results revealed that RTs were slowest following 4.5 Hz rhythms compared to faster (6.5 Hz) or slower (2.5 Hz) rates. It is possible that the 4.5 Hz rate similarity between non-speech woodblock cues and target double-vowel mixtures elicited more perceptual confusion (i.e., cognitive dissonance) at the “ideal speech rate” 4.5 Hz condition. Alternatively, 2.5 and 6.5 Hz may have elicited equally faster RTs according to a typical arousal/fatigue model of performance (Yerkes and Dodson, 1908). Either way, our results suggest a difference in rhythmic cue processing at 4.5 Hz that warrants further investigation.

We also observed an interaction between phase and rhythm type; within the PP rhythm condition, RTs were slowest for the early (-90°) phase condition. Thus, for perfectly periodic rhythms, the early phase (where targets occurred a quarter-interval before their expected position) was the most disruptive to subsequent speech perception. In contrast, the 180° phase could be perceived as a late offbeat, which is a defining characteristic of many Western popular music styles and therefore may not be as disruptive (Baur, 2021). That this interaction occurred for the PP rhythm condition suggests that this rhythm established the most robust temporal expectation through its high periodicity and predictability, perhaps explaining why this condition demonstrated greater RT disruptions than AP/AU conditions. The magnitude of RT improvement we find with forward entrainment is consistent with prior studies which similarly show a 40-100 ms change in RTs (Lange, 2009; Ellis and Jones, 2010; Rimmele et al., 2011). Several studies have similarly shown effects of forward entrainment in RT paradigms with or without changes in perceptual accuracy (reviewed by Saberi and Hickok, 2023).

### 4.3 Musical pulse continuation abilities relate to music training

Better musical pulse continuation abilities (as measured by the BDAT) predicted better speech identification accuracy in our entrainment task (**Fig. 2B**). This finding suggests the presence of domain-general entrainment abilities which support the perceptual continuation of temporal information from musical beats and rhythmic cues (Rammsayer et al., 2012). Such entrainment may operate via oscillatory mechanisms which form a type of predictive coding (Doelling and Poeppel, 2015). The explanatory power of BDAT indicates that individuals with a strong internal musical pulse performed better in rhythmically-cued speech perception. As our task did not contain a condition without rhythmic cueing, our data cannot answer whether generalized pulse continuation abilities explain concurrent speech perception more broadly. Still, links between atypical rhythm abilities and developmental deficits in speech-language disorders support general associations between auditory temporal processing and receptive speech communication (Vandermosten et al., 2010; Goswami, 2011; Ladányi et al., 2020).

We also replicate prior studies demonstrating that musical pulse continuation performance correlates positively with music training (Cinelyte et al., 2022). Listeners with more self-reported music training showed better internal mental representation of the musical beat (**Fig. 4A**). This result adds to a large body of evidence suggesting stronger auditory perceptual skills (e.g., Parbery-Clark et al., 2009) and improved temporal processing (Price and Bidelman, 2022; Perron et al., 2026) in trained musicians. As music training did not correlate with task accuracy, it seems that BDAT temporal skills predict aspects of speech perception *independently* of formal musicianship. It remains to be seen whether internal pulse or neural entrainment could be specifically trained to benefit SIN perception, though future longitudinal studies could address this question.

### 4.4 No relation between entrainment measures and SIN performance

Based on prior work (Bidelman and Yellamsetty, 2017), we anticipated that double vowel identification performance would correlate with SIN performance as measured by the QuickSIN (Killion et al., 2004). Though trends were in the expected direction, QuickSIN scores did not correlate with either BDAT pulse continuation thresholds or entrainment task performance. This suggests our cued speech task relies on mechanisms distinct from the perception of continuous speech. Additionally, it is entirely possible and even likely that rhythmic cueing may have a much stronger effect on the perception of continuous speech under noise degradation and at near threshold levels where other faciliatory mechanisms of signal-in-noise enhancement can take effect (McAuley et al., 2020; Shukla and Bidelman, 2021) (e.g., stochastic resonance). Future studies are needed to test this possibility.

## 5 Conflict of Interest

The authors declare that the research was conducted in the absence of any commercial or financial relationships that could be construed as a potential conflict of interest.

## 6 Author Contributions

JM: Conceptualization, Data curation, Formal analysis, Investigation, Methodology, Project administration, Resources, Software, Supervision, Validation, Visualization, Writing – original draft, Writing – review & editing; MZ: Investigation, Resources, Software, Writing – review & editing; GB: Conceptualization, Data curation, Formal analysis, Funding acquisition, Methodology, Resources, Software, Supervision, Visualization, Writing – original draft, Writing – review & editing

## 7 Acknowledgments

The authors thank Rose Rizzi and Jack Stirn for their feedback on early versions of this project and the study participants for their efforts and time.

## Data Availability Statement

The data for this study are available upon request to the corresponding author (G.M.B.).

## Notes

### Competing Interest Statement

The authors have declared no competing interest.

